# Endurance training restores ageing-impaired lysosomal biogenesis factors in rest and response to acute exercise in rat skeletal muscle

**DOI:** 10.1101/2025.01.28.635321

**Authors:** Hamid Rajabi, Benyamin Askari, David C. Clarke

**Author notes:** Corresponding author for correspondence contact Benyamin Askari.

## Abstract

**Purpose:** Lysosomes, crucial for autophagy, play a pivotal role in cellular processes influenced by exercise. This study investigates the impact of ageing on lysosomal function, focusing on Transcription Factor E3 and its regulators, mTORC1 and Calcineurin, emphasizing their response and adaptation to endurance training

**Methods:** Twenty-five male Sprague-Dawley rats were categorised into Young (2 months), Aged sedentary and Aged sedentary single session (18 months), Aged long-term trained, and Aged trained-single session (17 months). Changes in variables were explored concerning sarcopenia by Soleus muscle fibre diameter and number measured via Eosin & Hematoxylin, phosphorylated and total TFE3 protein levels via western blot, mTORC1 and Calcineurin mRNA levels via real-time PCR.

**Results:** Ageing occurred with increased pTFE3/TFE3_total_ protein (BF=579), declined mTORC1 mRNA (BF=3.99), and muscle diameter (BF=87), signifying sarcopenia and potential contributors. Conversely, Calcineurin mRNA (BF=0.67) and muscle fibre number (BF=0.31) remained unaltered during ageing. Exercise elicited acute responses, reducing pTFE3/TFE3_total_ protein (BF=306), elevating mTORC1 (BF=1.57) and Calcineurin mRNA levels (BF=3.19). Three weeks of endurance training further decreased resting pTFE3/TFE3_total_ protein (BF=174) while increasing Calcineurin mRNA (BF=12) and muscle fibre diameter (BF=126), with no changes in mTORC1 mRNA levels (BF=0.46) and muscle fibre number. Post-exercise, trained rats sustained decreased pTFE3/TFE3_total_ protein (BF=56) and elevated mTORC1 mRNA (BF=1.16).

**Conclusion:** This study underscores the involvement of TFE3, MTORC1, and Calcineurin in sarcopenia, proposing endurance training as an effective strategy to mitigate age-related changes and enhance muscle function in aged rats. Additionally, it suggests impairments in TFE3 possibly contribute to sarcopenia.

**key point:**

- Given the pivotal role lysosomes play in multiple homeostatic processes, investigating mTORC1, Calcineurin, and TFE3, an overlooked lysosome biogenesis factor involved in the metabolic effects of exercise, could help understand the metabolic state of sarcopenia and the role exercise plays.
- Through eosin & hematoxylin, western blotting and real-time PCR, we found Ageing results in sarcopenia, reduced TFE3 activity and mTORC1 gene expression.
- We saw a single bout of Endurance training elicited a response by increasing TFE3 protein activation, mTORC1, and Calcineurin gene expression, which is directed to improved sarcopenia.
- Three weeks of endurance training improved sarcopenia and was accompanied by elevated resting levels of TFE3 protein activation, and Calcineurin gene expression.
- Endurance training was still able to elicit post exercise response in TFE3 protein activation, and Calcineurin gene expression.
- Endurance training is a beneficial for sarcopenia, and TFE3 protein is a major player in inducing its effects.

## Introduction

Sarcopenia is characterized by the loss of muscle mass, strength, and performance that occurs with ageing and reduces exercise capacity. Sarcopenia is one of the leading causes of frailty syndrome in old age, which increases the risk of complications such as falls, fractures, physical disability, insulin resistance, diabetes, cardiovascular disease, and mortality(Bilski et al., 2022).

Among the contributing mechanisms to sarcopenia, protein and organelle homeostasis act as a mediator in muscle phenotypic changes, and the autophagy-lysosome system is a critical member of them(Triolo & Hood, 2021) capable of inhibiting cellular senescence(Carosi et al., 2022). Autophagy flux increases with age until the stage of lysosome transfer(Triolo & Hood, 2021). As a result, it is critical to focus on lysosome efficiency in the elderly because the available data on autophagic activity in aged muscle is conflicting. In old age, there is evidence of both increased and declined autophagy activity. Studies documenting excessive autophagy activity in old age, demonstrate that it results in further protein loss. Therefore, both insufficient and excessive autophagy causes muscle weakness, degeneration and atrophy respectively(Liang et al., 2020).

Lysosome dysfunction significantly contributes to autophagy-lysosome system inefficiency. However, regarding ageing, researchers have primarily studied autophagy-related proteins upstream of the lysosomes, and there is little information about its changes in young and aged muscle(Triolo & Hood, 2021).

Lysosomes are acidic organelles that play an important role in the autophagy and endocytosis systems. Lysosomes break down proteins and lipids, regulate cholesterol(Wróbel et al., 2022) and aid in the regeneration of the cell membrane in the event of damage(Corrotte & Castro-Gomes, 2019). Furthermore, they function as messenger organelles, creating specific pathways to control metabolism in response to any inefficiency or reduction in cell nutrients(Wróbel et al., 2022).

Transcription Factor E3 (TFE3) is one of the microphthalmia family of transcription factors (MiT-TFE) controling the gene expression of autophagy-lysosome system components(Triolo & Hood, 2021). When activated, these transcription factors translocate from the cytosol to the nucleus and promote the transcription of genes whose products participate in lysosome biogenesis. When lysosomes are healthy, and nutrients are abundant, mechanistic target of rapamycin complex 1 (mTORC1) kinase is activated by recruitment of microphthalmia family of transcription factors (MiT-TFE) to the lysosomes, then mTORC1 inhibits TFE3 by phosphorylating it and preventing its nuclear translocation. On the other hand, lysosomes inhibit mTORC1-specific phosphorylation in the absence of nutrients, demonstrating the lysosomes’ two-way interactions with mTORC1(Carosi et al., 2022; Wróbel et al., 2022). According to some evidence, mTORC1 activity in the aged muscle remains elevated and increases the severity of atrophy(Triolo & Hood, 2021). Accordingly, inhibiting mTORC1 slows biological ageing and improves health(Carosi et al., 2022).

Dysfunctional lysosomes release calcium ions via channels known as Mucolipin-1 protein (MCOLN1) which subsequently activates Calcineurin. Next, Calcineurin which is a Calcium ion and a Calmodulin-dependent protein phosphatase facilitates TFE3 nuclear translocation by dephosphorylating it(Wróbel et al., 2022). TFE3 then promotes autophagy and lysosome function via the coordinated lysosomal expression and regulation network(Triolo & Hood, 2021). It should be noted that the role of calcium ion signalling in TFE3 regulation is complex, with some evidence indicating that lysosomal calcium increases mTORC1 activity, which inhibits MiT-TFE signalling(Wróbel et al., 2022). Furthermore, given the increase in calcium-ion and free radicals in old age and their known relationship with TFE3, studying this factor in ageing can help with sarcopenia treatment. Few studies have been conducted in this field, with the majority focusing on genes and enzymes(Triolo & Hood, 2021).

By attenuating the age-related deterioration of skeletal muscle health, exercise emerges as a highly potent intervention that enhances not only muscle function and quality of life but also prolongs lifespan(Wang et al., 2022). Regular bouts of exercise provide a plethora of metabolic benefits in a variety of tissues, but even a single bout of exercise can initiate the activation of molecular pathways and adaptations important to health(Roberts & Markby, 2021). Exercise can activate autophagy and regulate its function. Exercise-induced autophagy especially by endurance exercise, is the most effective treatment for delaying sarcopenia and improving mitophagy, mitochondrial quality, and the number of inactive satellite cells, all of which rely on basal autophagy(Liang et al., 2020; Memme et al., 2021). It also promotes the degradation of dysfunctional proteins in skeletal muscle, produces energy substrates for skeletal muscle contraction, and suppresses apoptosis signals(Liang et al., 2020; Wang et al., 2022). Moreover, chronic exercise elevates the expression of several lysosomal markers, such as MCOLN1, which induces lysosomal biogenesis and enhances the muscle’s capacity for organelle turnover(Memme et al., 2021; Oliveira et al., 2021; Oliveira & Hood, 2019). Furthermore, in response to acute exercise, mTORC1, a known autophagy suppressor, is inhibited(Memme et al., 2021; Oliveira & Hood, 2019; Wang et al., 2022). Meanwhile, transient Ca2+ fluxes increase and activate Calcineurin(Memme et al., 2021; Oliveira & Hood, 2019). Together, they mediate TFE3’s dephosphorylation and its nuclear translocation(Memme et al., 2021; Oliveira & Hood, 2019; Roberts & Markby, 2021). Finally, after entering the nucleus, TFE3 controls several lysosomal and autophagy-related genes(Oliveira & Hood, 2019).

While numerous studies have investigated the effects of endurance training on skeletal muscle and autophagy, limited attention has been given to lysosomes and their associated factors, particularly their response following exercise(Baek et al., 2022). Given the crucial role of lysosomes in maintaining cellular homeostasis, the established exercise benefits, and TFE3’s role in the metabolic effects of exercise, further investigation is needed to understand the factors contributing to lysosome biogenesis in young and elderly, as well as how exercise training, affects them at rest and post-exercise(Roberts & Markby, 2021). Therefore, the aims of this study were as follows: 1. To assess the occurrence and extent of sarcopenia concerning ageing by evaluating Soleus muscle fibre diameter and number. 2. To evaluate lysosome function by measuring the protein level of pTFE3/TFE3total, as the primary factor, along with the mRNA levels of two upstream factors, mTORC1 and Calcineurin, in different conditions such as youth, ageing, long-term exercise, their post-exercise responses, and effect of training adaptation on their post-exercise response magnitude. 3. To investigate the impact of lysosome biogenesis factors on muscle fibre characteristics, establishing any direct relationships between these factors and muscle fibre characteristics.

## Methods

### Ethical approval

The Ethics Committee of the Research Institute of Sports Sciences, Ministry of Science, approved the study **(IR.SSRI.REC.1399.760)**. Rats were purchased from the Pasteur Institute in Tehran, Iran, and five animals were housed in each cage at 22°C, 60% humidity, and a light-dark cycle of 12:12 hours with Ad libitum feeding. Animal handling and anaesthetization followed American Veterinary Association guidelines(Leary et al., 2020) and “Principles of laboratory animal care”(National Research Council (US) Committee for the Update of the Guide for the

Care and Use of Laboratory Animals, 2011). Each rat was individually isolated and anaesthetized with carbon dioxide one by one, followed by decapitation, and their tissues were sampled. The corresponding author received training in animal surgery and handling prior to the study through courses at the Sports Science Research Institute and Pasteur Institute.

### Animals

A total of 25 male Sprague Dawley rats at three age points were assigned into five groups of five animals per cage: young (2 months, 244±9.5 g), aged sedentary (A-SED) (18 months, 395±9.2 g), aged sedentary single session (A-SEDS)(18 months, 403±7.6 g), aged long-term trained (A-LT)(17 months, 356±23.2 g), and aged trained-single session (A-TS)(17 months, 358±17.7 g) to control for effects of ageing, training adaptation and acute response. The sample size was determined using a previously proposed formula(Charan & Kantharia, 2013). In this formula, “E” is a degree of freedom ranging from 10 to 20. If E is less than 10, the likelihood of a false negative increases, but any value greater than 20 does not affect the likelihood of a significant result. Although “E” is based on the Analysis of variance (ANOVA) test, it applies to all statistical tests and does not affect the results. Based on the mentioned formula the sample size was determined to be 25 which is optimal.

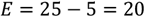

All animals were weighed at the beginning of the project. The two trained groups were also weighed after completing their respective training protocol. The animals’ species, sex, training protocol, and related procedures followed a previously published protocol (Zhu et al., 2016). The corresponding author performed all procedures, including animal handling, surgeries, experimental and data analyses. No blinding was implemented at any stage of the experiment, and the author was aware of the group allocations throughout the study.

### Exercise intervention

A motor-driven treadmill with eight lines (Tajhiz Gostar Omid Iran, Iran) was used for the exercise training. Rats in the exercise groups first underwent three days of familiarization prior to the experiment. During this period, the rats walked on a treadmill at 10 m/min for 20 min/day on a 0° incline. In the experiment, rats willing to undergo training (rats subjected to electric shock [1.0 mA] 5 times, with each shock lasting 1s) were used. All rats in this study met the required conditions. The training protocol for each session was 15 m/min for 30 min/day on a 0° incline. The intensity of the shock stimulus was set to 1.5 mA. Following the familiarization phase, the A-SEDS group did one session of treadmill running at 15 m/min for 30 min at a 0° incline. 3h after exercise, this group, along with the young and A-SED groups, were euthanized. During the experiment, the A-LT group and the A-TS groups were subjected to three weeks of 6-days per week treadmill training at 15 m/min, 30 min/d at 0° incline. The A-TS group and the A-LT group were euthanized 3h and 24h after exercise respectively. Both groups were weighed again before the euthanization.

### Tissue processing and molding

After separating the sample tissue of the soleus muscle from the right leg, it was immediately placed in 10% formalin solution for 24 hours followed by dewatering using ethyl-alcohol (70%, 80%, 90%, 100%) in four, 50-minute stages respectively. Next, the alcohol was removed and the sample was prepared for molding and cutting by incubating the tissues in two xylol and two paraffin changes for 50 minutes each respectively. Muscle samples were sectioned using a microtome set at 5-μm thickness and then placed on a silanized slide.

### Hematoxylin & Eosin staining

After melting (Vinteb) the sample’s paraffin the slides were placed in two Xylol changes (Sigma-1330-20-7) for 15 minutes each. Prior to staining, the tissue was hydrated through alcohol solutions (100%, 90%, 80%, 70%) and distilled water for 5 minutes each respectively. Next, slides were placed in hematoxylin dye (Sigma-H9627) for 7 seconds, then washed for 1 minute in distilled water. They were placed in lithium carbonate (Sigma-1.05680) for 2 seconds and then in Eosin dye (Sigma-HT110116) for 3 minutes. Afterward, the sample was dehydrated by incubation in alcohol solutions (90%, 100%, 100%) for 4 seconds each respectively. Then, for clarification, the sample was placed inside two Xylol changes for 15 minutes each. In the end, a drop of Entlan glue (Sigma-1.07961) was placed on the sample and the coverslip was glued on the slide to get photographed with an optical microscope (LABOMED).

### Western blotting

For total TFE3 protein Lysed tissue samples were separated by SDS-PAGE (Sigma-Aldrich, Germany) and the antibody was (Cell Signaling Technology Cat# 14779, RRID: AB_2687582). The separated proteins were transferred to the PVDF membrane (Sigma-Aldrich, USA) with 0.45-micrometre pores, then proteins were transferred according to the manufacturer’s instructions. At room temperature, the PVDF paper floated in the blocker solution and placed on a shaker for 2 hours before getting washed with TBST. Next, the PVDF paper in the primary antibody was kept overnight in the refrigerator. Then, it was washed with TBST 3 times and the PVDF membrane was placed in the secondary antibody solution for another 2 hours on a shaker at room temperature before getting washed 3 times again with TBST. Finally, two ECL kit solutions (Pars tous, Iran), X-ray film (Fujifilm, Japan) and ImageJ (RRID:SCR_003070) software used for revealing and analysing the bands. For phosphorylated TFE3, the same procedure as (Martina et al., 2016) was used.

### cDNA synthesis and RNA extraction

Kit materials (Parstous, Iran) and RNA samples were first vortexed (KIAGENE, Iran) and spun then, RT buffer, RT enzyme, Oligo dT primer, and DEPC water were mixed to make cDNA for RT mix. Next, the mix was distributed in 9 μl volumes and was poured into 0.2 ml microtubes. At last, the RT mix and RNA sample were placed in a dry block heater (KIAGENE, Iran) and the temperature program was executed (25°C - 10min, 47°C - 60min, 85°C – 5min, 4°C – 2min). The tissue sample was first frozen in liquid nitrogen and powdered in a crucible, then 1 ml of Trizol (Kiazist, Iran) was added for every 10 million isolated cells. 200 microliters of chloroform (DRM-CHEM, Iran) were added to the samples per 1 ml of trizol, then shaken for 15 seconds without vortex. The microtubes were incubated for 15 minutes at room temperature, then centrifuged (Hettich, Germany) for another 15 minutes at 12000 RPM and 4°C. Next, the clear supernatant part containing RNA was transferred to another microtube got mixed with isopropanol (DRM-CHEM, Iran). In the next step, 500 microliters of trizol were added per 1 ml of the solution and were mixed by inverting before getting incubated for 10 minutes at (−20°C) temperature. Then, the samples were centrifuged for 10 minutes at 12000 RPM, 4°C, and the sediments were washed with 75% ethanol after draining the supernatant solution. Afterward, the solution was centrifuged for another 5 minutes at 7500 RPM, 4°C. Finally, the supernatant was drained again and the microtubes were placed on a sterile paper towel for 15 minutes to sediment dry and the alcohol evaporates.

### Realtime-PCR

First, all the necessary materials for PCR were vortexed (KIAGENE, Iran) and spun. Next, for each gene, a mixture of PCR components PCR mix (addbio, Korea), reverse and forward primer (sinaclon, Iran), and water were mixed and spun before distributing in device microtubes. 1μl cDNA sample was added to each vial (the final volume of each PCR reaction is 10 μl) then vortexed and mixed again before execution of the temperature program (ABI Stepone, USA). At last, the CT numbers of the reference and the main genes were calculated in the 2^-DCT^ formula, for relative gene expression changes. The primer design for Calcineurin and mTORC1 mRNA is provided in (Table 1).

**Table 1.**
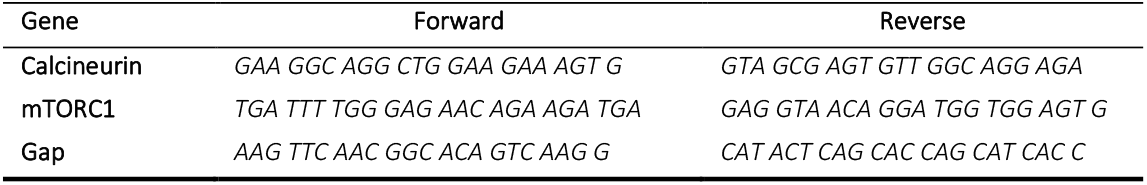
Primer design for Calcineurin and mTORC1 mRNA.

### Statistical analysis

Because of some shortcomings of the “p-value” and proposed solutions (Wasserstein et al., 2019), Bayesian statistics were used for analysis. For a better understanding of the results, “p-value” and effect sizes are also reported. JASP software (Version 0.16. 2) (RRID:SCR_015823) was used to preprocess, analyze, and visualize the data. The distribution of data was assessed using the Shapiro-Wilk test to determine the appropriateness of subsequent statistical analyses. First, the Bayesian Pearson/Kendall correlation coefficient and Regression analyses were used to assess the relationships between variables. Depending on the number of groups and the hypothesis, independent t-test, paired sample t-test, or one-way ANOVA followed by the Games-Howell post-hoc test were used. The analysis results are reported under the Bayesian Analysis Reporting Guidelines (Kruschke, 2021). All data points included in the analysis.

## Results

In addition to analysing and comparing the results between groups and assessing the post-training session response to a treadmill running exercise and the effect of training adaptation on the magnitude of this response, the values for pTFE3/Total TFE3, mTORC1, and Calcineurin of the A-SEDS were subtracted from the A-SED group, and A-TS subtracted from the A-LT group

### Body mass in young, aged sedentary and trained

The statistical test revealed that exercise did not affect the body mass in A-TS and A-LT groups: the reduction was not reliable (A-LT, anecdotal evidence, BF=0.966) (A-TS, anecdotal evidence, BF=0.744). As it is evident from (Figure 1), when the body mass of the A-SED group and the young group were compared, it was evident that the A-SED group had 61.58% more body mass (Decisive evidence, BF=372482). However, the A-LT group weighed 16% less than the A-SED group (Decisive evidence, BF=789) and 36% more than the young group (decisive evidence, BF=7631).

**Figure 1.**
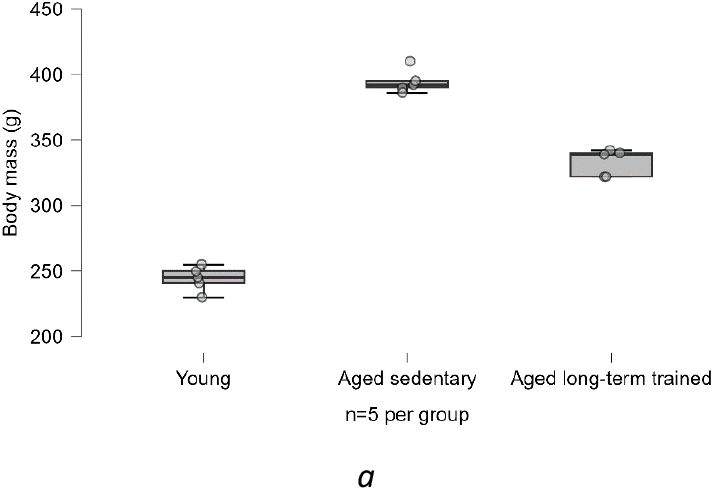
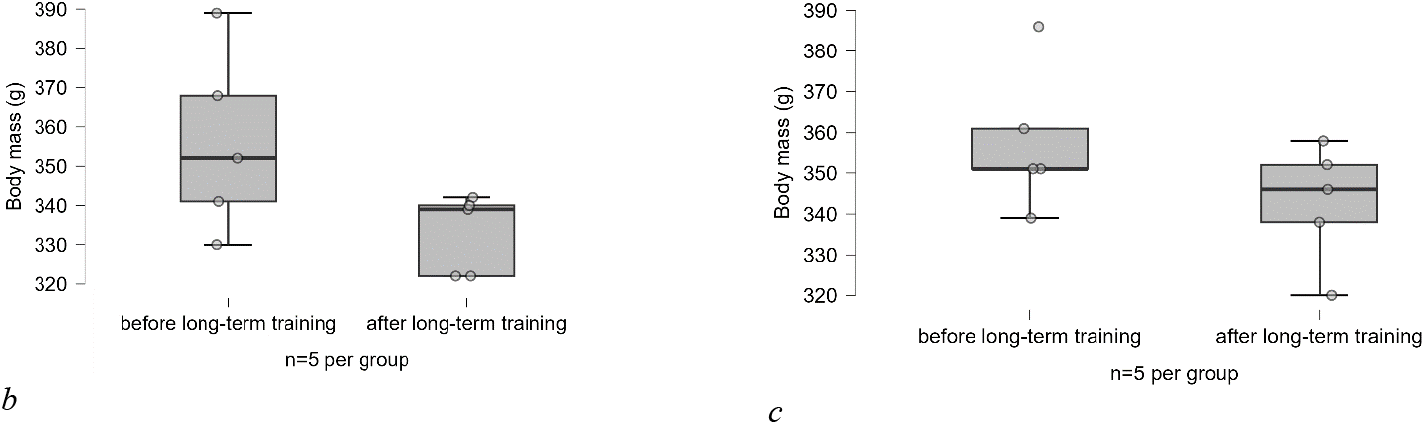
**a)**The body mass differences between the A-SED group with the young group (150g), (BF=372482), (p<0.001, Cohen’s d=-15.63); the A-LT group with the A-SED group (−61.6g), (BF=789), (p<0.001, Cohen’s d=6.4); and the A-LT group with the young group (88.8g), (BF=7631), (p<0.001, Cohen’s d=-9.23); **b)**The body mass reduction in the A-LT group (23 g), (BF=0.966), (p = 0.16, Cohen’s d = 0.75)**; c)**The A-TS group (14.8g), (BF=0.744), (p=0.24, Cohen’s d=0.6) after training

### Soleus muscle fibre diameter and number

As (Figure **2**) has demonstrated, with increasing age, the soleus muscle fibre diameter had decreased by 26.46% (very strong evidence, BF=87) when comparing A-SED with the young group, but regarding muscle fibre number no changes were observed (moderate evidence, BF=0.319). Comparing A-SED with A-LT, the trained group shows higher values both in muscle fibre diameter, 25.26% (Decisive evidence, BF=126) and muscle fibre number, 49% (very strong evidence, BF=70). In A-LT the muscle fibre number was 38.59% more than the young group (Figure **2**) (moderate evidence, BF=3.123). However, further explanations and comparisons regarding hypertrophy and hyperplasia are made in the discussion section.

**Figure 2.**
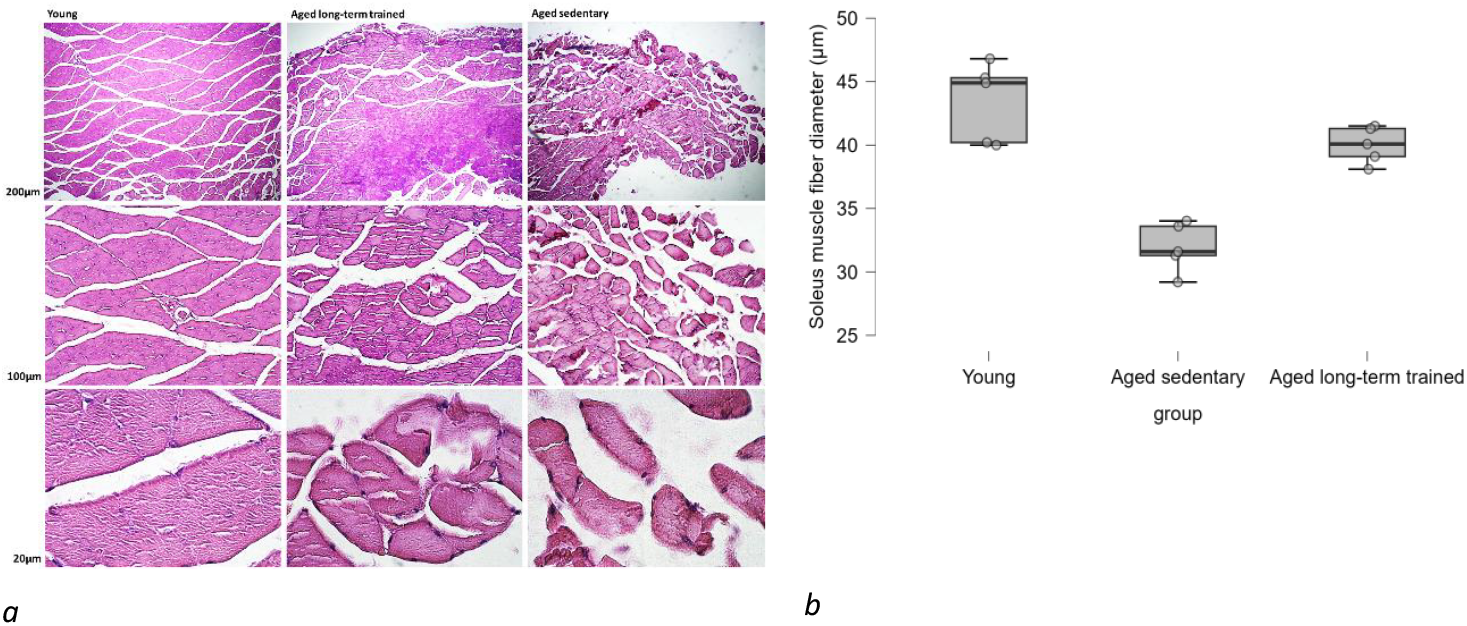
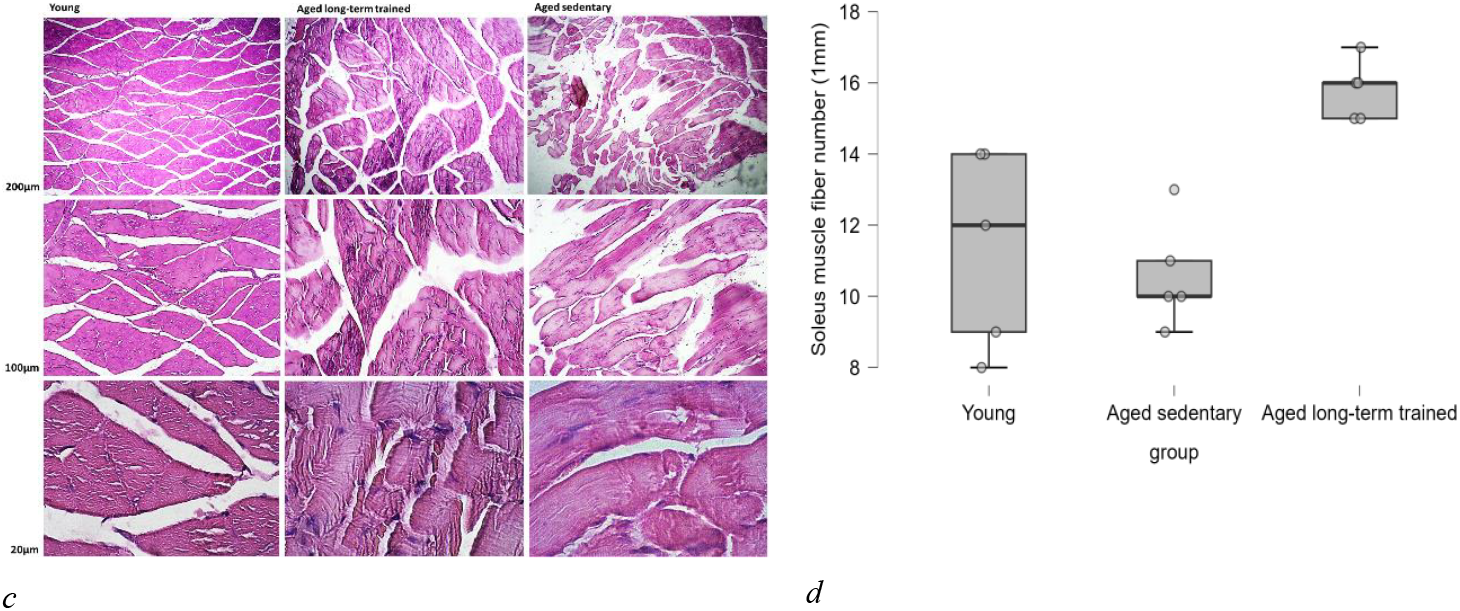
Changes in soleus muscle fibre diameter and number through ageing and by training measured by Hematoxylin & Eosin staining. **a)**and **b)**Differences in the soleus muscle fibre diameter between the A-SED group and the young group (11.49 µm), (BF=87), (p<0.001, Cohen’s d=4.41); between the A-LT group and the A-SED group (8.07 µm), (BF=126), (p<0.001, Cohen’s d=-4.71); and between the A-LT group and young (3.42 µm), (BF=1), (p=0.15, Cohen’s d=1.4). **c)**and **d)**Differences in the soleus muscle fibre number between the A-SED group and the young group (0.8 u), (BF=0.319), (p=0.8, Cohen’s d=0.35); between the A-LT group and the A-SED group (5.2 u),(BF=70),(p<0.001, Cohen’s d=-4.24); and between the A-LT group and the young group (4.4 u) (BF=3.123), (p=0.04, Cohen’s d=-2.13).

### TFE3 protein levels

The variable pTFE3/TFE3total is calculated as a measure of this protein’s activity in the muscle; the lower the phosphorylated (inactive type of this protein compared to its total expression) the more active it is. In the comparison of young and A-SED groups, the aged group nearly had twice the values of pTFE3/Total TFE3 (195% increase) (Decisive evidence, BF=579) demonstrating the effects of ageing on TFE3 protein activity (Figure **3**). Regarding the post-exercise response of this protein to a single bout of treadmill running, as Figure3 has shown, the A-SEDS group had 54.59% less pTFE3/Total TFE3 than the A-SED group (Decisive evidence, BF=306). After three weeks of endurance training, the A-LT group had 56.75% less pTFE3/Total TFE3 than the A-SED group (Decisive evidence, BF=174) showing at the baseline level, the inactive TFE3 has declined while still being 27.65% higher than the young group (Moderate evidence, BF=7). As visualized in (Figure **3**), when the response of this protein post-exercise session was evaluated after three weeks of endurance training, it was discovered that the aged single bout after the long-term training group had 31.7% lower pTFE3/Total TFE3 values immediately after post running session compared to the aged long-term training group (very strong evidence, BF=56) showing that even after three weeks of training, endurance exercise still elicited post session response of TFE3.

**Figure 3.**
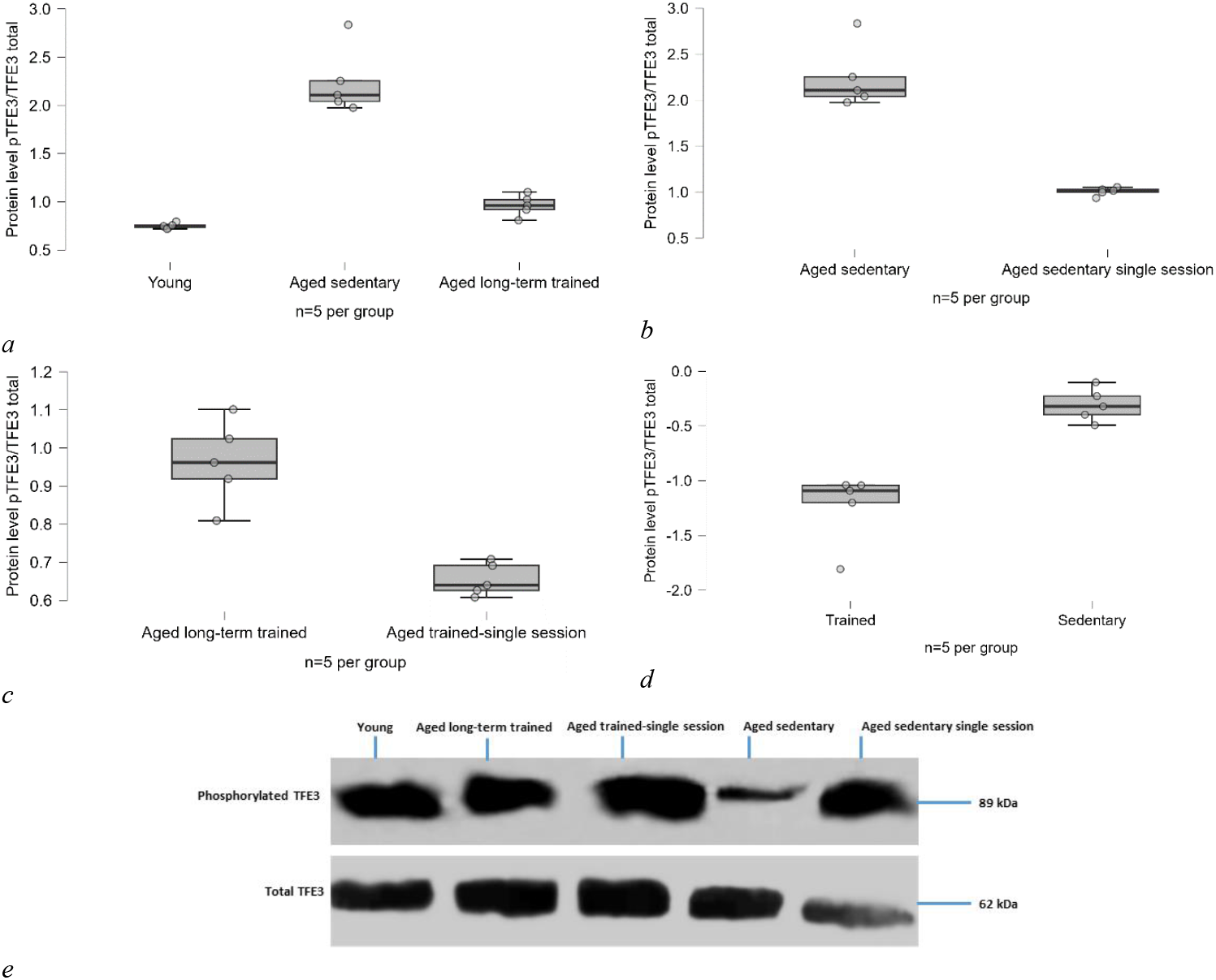
**a)**Differences in the soleus muscle pTFE3/TFE3_total_ at rest between the A-SED group and the young group (1.462 u), (BF=579), (p<0.001, Cohen’s d=-7.08) and the A-LT group (1.255 u), (BF=174), (p<0.001, Cohen’s d=6.07); between the A-LT group and young group (0.207 u), (BF=7), (p=0.02, Cohen’s d=-1.01). **b)**Differences in the soleus muscle pTFE3/TFE3_total_ in response to a single bout of exercise compared between the A-SEDS group and the A-SED group (1.207 u), (BF=306), (p<0.001, Cohen’s d=5); **c)**between the A-TS group and the A-LT (0.303 u), (BF=56), (p<0.001, Cohen’s d=3.69). **d)**Differences in the magnitude of the soleus muscle pTFE3/TFE3_total_ response to a single bout of exercise compared between sedentary and trained groups (0.904 u), (BF=53), (p<0.001, Cohen’s d=-3.65). **e)**Image of the Western Blotting Band for the results of skeletal muscle synthesis-related protein analysis at rest and response to exercise.

Between the aged sedentary and aged trained groups, the trained group’s post-session pTFE3/Total TFE3 response magnitude was 74.88% lower than the sedentary group (very strong evidence, BF=53) (Figure **3**).

### mTORC1 mRNA levels

The A-SED group had 52.89% less mTORC1 mRNA levels than the young group (moderate evidence, BF=3.99) (Figure 4). In terms of post-session response to endurance exercise, the A-SEDS group had 86.99% more mTORC1 mRNA levels than the A-SED group (Anecdotal evidence, BF=1.57). However, the mTORC1 mRNA levels in the A-LT group compared to the A-SED group did not change (anecdotal evidence, BF=0.462) (Figure 4). Further, despite showing 70.58% more mTORC1 mRNA levels than the A-LT group, the results for the response of mTORC1 mRNA levels to a single bout of exercise in the A-TS group suggests a small probability of post-exercise response (anecdotal evidence, BF=1.168). Moreover, the magnitude of the changes in mTORC1 mRNA after three weeks of treadmill running did not show significant changes when comparing sedentary and trained groups (anecdotal evidence, BF=0.558).

**Figure 4.**
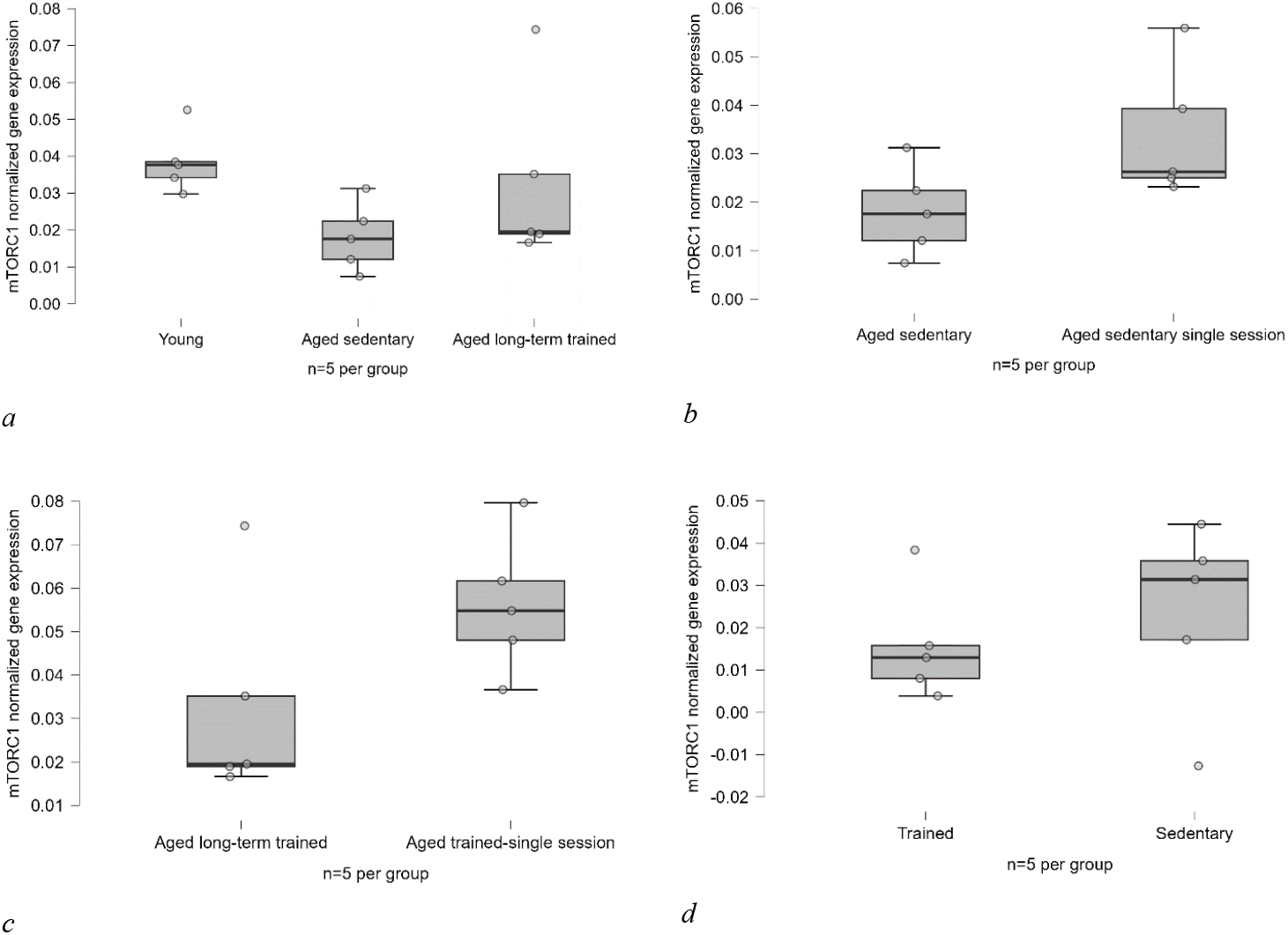
**a)**Differences in the soleus muscle mTORC1 mRNA at rest between the A-SED group and both the young group (0.02 u), (BF=3.99), (p=0.017, Cohen’s d=1.29) and the A-LT group (0.0014 u), (BF=0.462), (p=0.46, Cohen’s d=0.93); between the A-LT group and young group (0.005 u), (BF=0.311), (p=0.88, Cohen’s d=0.35). **b)**Differences of the soleus muscle mTORC1 mRNA in response to a single bout of exercise compared between the A-SEDS group and the A-SED group (0.015 u), (BF=1.57), (p=0.06, Cohen’s d=-1.34); **e)**between the A-TS group and the A-LT (0.023 u), (BF=1.168), (p=0.11, Cohen’s d=-1.12). **d)**Differences in the magnitude of the soleus muscle mTORC1 mRNA response to a single bout of exercise compared between sedentary and trained groups (0.007 u), (BF=0.558), (p=0.54, Cohen’s d=-0.403).

### Calcineurin mRNA levels

The Calcineurin mRNA levels did not differ between young and A-SED group (anecdotal evidence, BF=0.675) (Figure 5). However, in response to a single bout of treadmill running exercise, Calcineurin mRNA levels was 85.11% higher in the A-SEDS group than in the A-SED group (moderate evidence, BF=3.197). Following three weeks of treadmill running, the A-LT group had three times 153.31% more Calcineurin mRNA levels than the A-SED group (strong evidence, BF=12).

**Figure 5.**
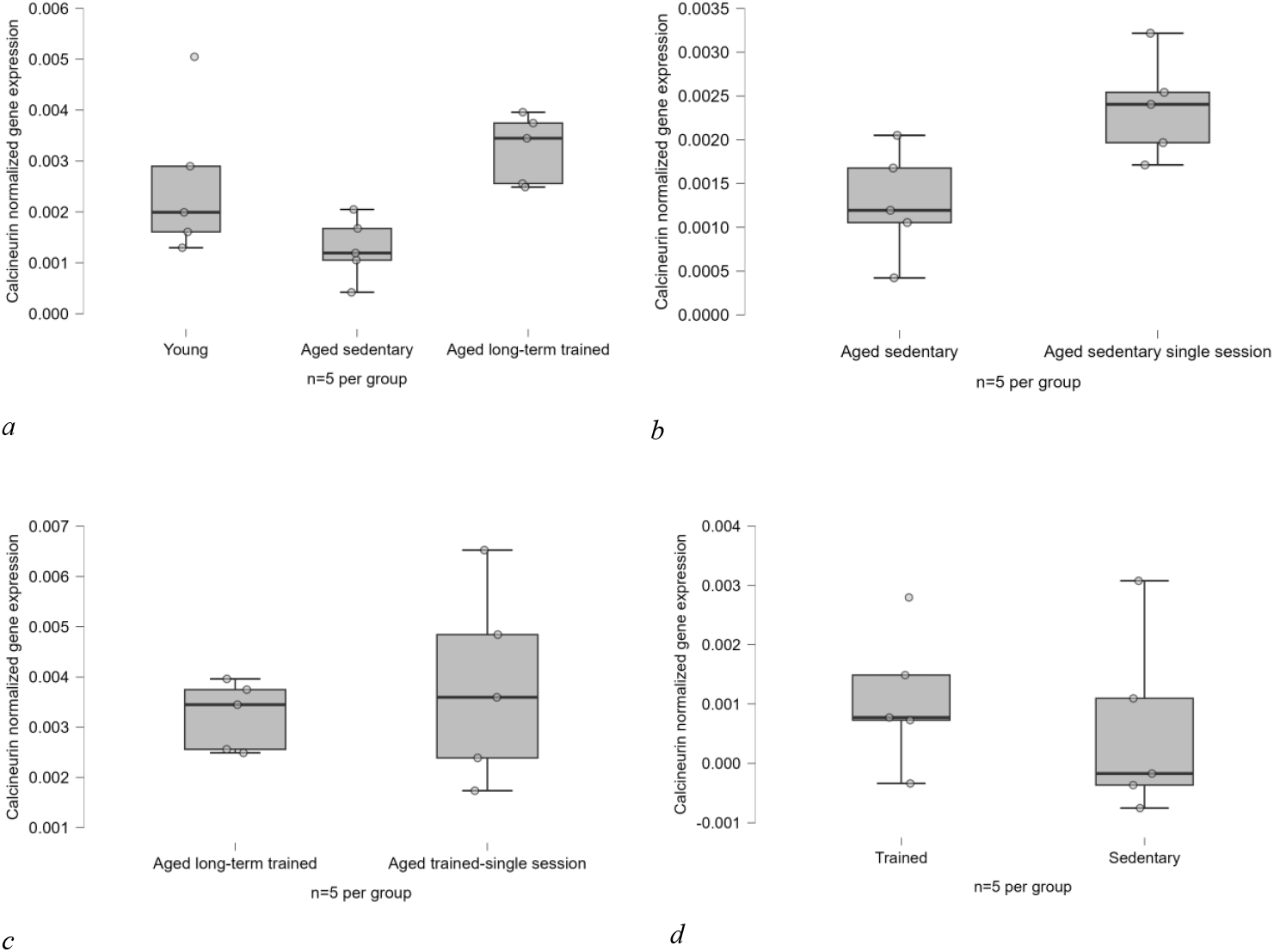
**a)**Differences in the soleus muscle Calcineurin mRNA at rest between the A-SED group with both the young group (0.0012 u), (BF=0.675), (p=0.26, Cohen’s d=1.11) and the Aged-LT group (0.0019 u), (BF=12), (p=0.004, Cohen’s d=3.01); and between the A-LT group with young group (0.0006 u), (BF=0.371), (p=0.65, Cohen’s d=0.57). **b)**Differences of the soleus muscle Calcineurin mRNA in response to a single bout of exercise compared between the A-SEDS group and the A-SED group (0.001 u), (BF=3.197), (p=0.02, Cohen’s d=-1.81); **c)**between the A-TS group and the A-LT (0.00057 u), (BF=0.557), (p=0.54, Cohen’s d=-0.39). **d)**Differences in the magnitude of the soleus muscle Calcineurin mRNA response to a single bout of exercise compared between sedentary and trained groups (0.00051 u), (BF=0.549), (p=0.021, Cohen’s d=1.81).

In contrast, Calcineurin mRNA levels in the A-TS group did not differ from that in the A-LT group (anecdotal evidence, BF=0.557) (Figure 5). Similarly, difference was observed or there was only a trend when comparing the effect of three weeks of treadmill running on the magnitude of post-exercise response between the sedentary and trained groups (anecdotal evidence, BF=0.549) (Figure 5).

### The relationship between variables

Based on Bayesian Kendall’s correlation test, except for pTFE3/total TFE3 and soleus muscle fibre diameter, there was no relationship between any of the measured variables. According to the Bayesian Kendall’s one-way correlation test, there was a strong reverse correlation between pTFE3/total TFE3 and soleus muscle fibre diameter, indicating that as phosphorylated TFE3 (inactive) increases, so does soleus muscle fibre diameter (β=-0.581, strong evidence, BF_-0_=42.557) (Table 2).

**Table 2.** Bayesian Kendall’s Tau Correlations.

|  | Kendall's tau B | BF<sub>0</sub> |
| --- | --- | --- |
| PTFE3 / TOTAL TFE3 - Soleus muscle fiber diameter ($\mu\text{m}$) | -0.581** | 42.557 |

\* $BF_0 > 10$ , \*\* $BF_0 > 30$ , \*\*\* $BF_0 > 100$
Note. For all tests, the alternative hypothesis specifies that the correlation is negative.

Following the correlation test, the Bayesian linear regression revealed pTFE3/total TFE3 accounts for 80% of the observed changes in soleus muscle fibre diameter (R^2^=0.809, decisive evidence, BF=2408.781), (Table 3).

**Table 3.** Model Comparison – Soleus muscle fibre diameter (µm)

| Models | P(M) | P(M\|data) | BF<sub>M</sub> | BF<sub>10</sub> | R<sup>2</sup> | Mean | Credible interval |  |
| --- | --- | --- | --- | --- | --- | --- | --- | --- |
|  |  |  |  |  |  |  | Lower | Upper |
| Null model | 0.500 | 4.150e-4 | 4.151e-4 | 1.000 | 0.000 | 38.467 | 37.075 | 39.751 |
| PTFE3 / TOTAL TFE3 | 0.500 | 1.000 | 2408.781 | 2408.781 | 0.809 | -6.538 | -8.522 | -4.712 |

Bayesian Linear Regression

## Discussion

This study aimed to examine the role of factors responsible for lysosome biogenesis in sarcopenia and then thoroughly evaluate their trainability by looking at how they adapt and respond to exercise and further, how the occurred adaptation affects their post-training response to fully decipher their dynamics.

### Factors regulating lysosome biogenesis and muscle fibre: effect of ageing and training

As expected, the soleus muscle fibre diameter in the aged sedentary group was smaller than the young group showing they had experienced sarcopenia which is aligned with another study observing similar results under similar conditions(Baek et al., 2022). The number of soleus muscle fibres, however, did not change with age (Figure **2**). On the metabolic side, pTFE3/TFE3total in the aged sedentary group increased to nearly three times that of the young group indicating a decrease in its activity and nuclear translocation, and thus reduced lysosome biogenesis, which explains the observed muscle wasting (Figure **3**).

Following the same trends, mTORC1 gene expression in the aged sedentary group was at nearly half of the values in the young group (Figure 4). In the context of ageing, the prevailing literature highlights prolonged mTORC1 activity, which is linked to autophagy inhibition and the subsequent development of sarcopenia. However, the existing body of evidence regarding autophagy presents a somewhat contradictory picture, with some studies indicating excessive autophagy and others suggesting under-activation, along with impairment of the autophagy/lysosome system. Next, due to the feedback-like relationship between mTORC1 with lysosomes(Yang et al., 2019), have reported that excessive mTORC1 inhibition causes enlarged lysosomes as well which exacerbates its dysfunction and is aligned with the observed decrease in TFE3 activity(Yang et al., 2019). In parallel with the findings of the current study, the observed decrease in mTORC1 and TFE3 activity implies a situation where ageing may be associated with excessive autophagy coupled with lysosome dysfunction(Liang et al., 2020; Triolo & Hood, 2021). Further, Calcineurin gene expression did not change with age from the statistical point of view (Bf=0.675), which can be assumed that the ca^2+^ remained unchanged in the aged sedentary due to being sedentary and subsequently, no changes in Calcineurin mRNA levels (Figure 5).

Three weeks of endurance training effectively improved sarcopenia and increased soleus muscle fibre diameter in the aged long-term training group, not only when compared to the aged sedentary group, but also as a trend in the samples, approaching that of the young group. (Figure **2**). Contrary to the unchanged soleus muscle fibre number in ageing, three weeks of endurance training led to an increase in the soleus muscle fibre number in the aged long-term training group compared to the aged sedentary group (Figure **2**). However, even though there is small evidence supporting the result of this study even in similar periods and sample age(Tamaki & Uchiyama, 1995), the available data on hyperplasia is elusive and the topic is rather debatable; therefore, the obtained results need to be put in perspective, and interpreted carefully to prevent false conclusions. The argument over the possibility of hyperplasia is not new with evidence both supporting and rejecting its occurrence as a result of training(Jorgenson et al., 2020). Nevertheless, considering the body of literature, it is unlikely that an actual increase in soleus muscle fibre number occurred as the result of endurance training because there is evidence suggesting most observed hyperplasia is a result of muscle architectural variables such as length of the fibre and/or its pennation angle (Jorgenson et al., 2020). Next, the aged long-term training group had lower resting pTFE3/total TFE3 values than the aged sedentary group, indicating that endurance training is effective in improving aged muscle via TFE3-induced lysosome biogenesis Figure **3**). However, the soleus muscle’s mTORC1 gene expression did not differ between the aged long-term training group and the aged sedentary group with three weeks of endurance training, which according to the literature could probably be due to changes in adenosine monophosphate-activated protein kinase (AMPK) caused by the endurance training and its regulatory effect on mTOR(Wang et al., 2022). Although evidence regarding the effect of endurance training on mTOR is somewhat complex with some showing its expression has increased after 12 weeks of 3-5 sessions of 60-minute exercise, others reported the inhibitory effect of endurance training on this factor(Wang et al., 2022) (Figure 4). Further, the higher resting Calcineurin gene expression in the aged long-term training group compared to the aged sedentary group (Figure 5), partly explains the increased TFE3 activity (Figure **3**) which leads to subsequent lysosome biogenesis and autophagy induction in the aged long-term training group and improved soleus muscle fibre diameter and number (Figure **2**). Although some evidence exists on the positive effect of Calcineurin on mTORC1(Wróbel et al., 2022; Yang et al., 2019), and mTORC1 on lysosome size, lysosome biogenesis is primarily inhibited by mTORC1 activation, and these relationships are regulatory and feedback-like, meaning they control lysosome function from excessive activity or inactivity thus it is not necessary to observe mTORC1 reinforcement followed by Calcineurin activity. It should also be noted that the results for mTORC1 activity (Figure 4) are completely aligned with another study which did not observe any difference between young sedentary, aged sedentary, and aged trained subjects. Although, in their study mTORC1 was examined at the protein level while in this research it was measured at the level of gene expression (Baek et al., 2022).

Regarding body mass changes, the aged long-term training and aged trained single session groups showed a slight decrease after completing the training protocol only as a sample trend (Figure 1). In contrast, when comparing the aged sedentary group to the aged long-term training group at the same age (18 months), the aged long-term training group had less body mass, demonstrating the effect of three weeks of endurance training on body mass (Figure 1). Based on the observed changes, it may be possible that the effect of endurance training on body mass in the aged long-term training and aged trained single session groups pre and post-training was neutralized by ageing as evidence suggests that both animal and human fat mass increases with age(Quirós Cognuck et al., 2020). The aged sedentary group had more body mass than the young group, possibly due to fat mass increase with age. However, this cannot be strongly concluded as fat mass was not measured. Furthermore, the higher body mass in the aged long-term training group compared to the young group suggests that three weeks of training did not fully compensate for the effects of sedentary ageing on body mass gain (Figure 1). Additionally, all the observations in the young, aged sedentary, and aged long-term training groups had previously been observed in another study(Baek et al., 2022).

Based on the observed strong causal effect of pTFE3/total TFE3 on the soleus muscle fibre diameter (R^2^=0.8) (Table 3) and the existing evidence about the effects of this protein and its upstream factors mTORC1 and Calcineurin on cellular processes and muscle breakdown, It could be assumed that the decrease in TFE3 activity (Figure **3**) by ageing leads to lysosome dysfunction and subsequently causes the cellular processes related to this organelle to function inefficiently, all of which explain the developed sarcopenia in the aged sedentary group (Figure **2**). However, to completely prove the causal relationship between TFE3 and soleus skeletal muscle fibre diameter, a methodological causal relationship evaluation is needed as well. According to another study (Roberts & Markby, 2021), TFE3 KO samples could not benefit from the beneficial effects of exercise. Therefore, put together with the statistical test in this study we could assume with more confidence that the TFE3 is responsible for the changes in soleus muscle fibre diameter.

### Effect of long-term training on the post-exercise response of factors regulating lysosome biogenesis and soleus muscle fibre diameter and number

A single bout of endurance exercise could elicit pTFE3/total TFE3 response in the aged single-session group compared to the aged sedentary group (Figure **3**). Moreover, the gene expression of mTORC1 (Figure 4) and Calcineurin (Figure 5) was also higher in the aged single session group than in the aged sedentary. The post-exercise session responses in TFE3 (Figure **3**) and Calcineurin (Figure 5) activity are aligned with the body of literature showing the anti-ageing effects of exercise begin from the first session, and that in response to a single bout of endurance exercise, increased Calcineurin dephosphorylates TFE3 while it could also explain some part of mTORC1 activity(Oliveira & Hood, 2019; Wróbel et al., 2022)(Figure 4). However, the results regarding mTORC1 are inconsistent with some studies reporting that mTORC1 is inhibited during and after exercise(Memme et al., 2021; Wang et al., 2022). This difference is not a contradiction in any way, since the increased mTORC1 could control the previously assumed excessive autophagy which is also consistent with mTORC1 expression in the aged sedentary group.

In the aged trained groups, the aged trained-single session group had less pTFE3/totalTFE3 and more mTORC1 gene expression post-training session compared to the aged long-term training group showing that still after three weeks of training, exercise elicited a post-session response (Figure **3**) (Figure 4). However, Calcineurin remained unchanged (Figure 5) which could be an indication of its adaptation to the exercise protocol. Furthermore, for a more accurate conclusion regarding post-exercise response and the effect of adaptation on its magnitude, we further examined these factors’ responses and compared them between sedentary and trained elderly groups. After three weeks of endurance training, the magnitude of the post-exercise response of pTFE3/totalTFE3 in the trained group was lower, but there was no significant difference in mTORC1 and Calcineurin gene expression, indicating that endurance training continued to have a regulatory effect on mTORC1 (Figure 4) (Figure 5) but pTFE3/totalTFE3 had adapted to some extent (Figure **3**). This additional analysis is useful for longer-term studies because it demonstrates that, to truly analyze the effect of exercise, progressive overload may be required in training protocols.

## CONCLUSION

Skeletal muscle atrophy with age and a considerable share of this atrophy is caused by a decrease in TFE3 protein activity, which is accompanied by a decrease in mTORC1 gene expression, implying the possibility of excessive autophagy activity associated with lysosome dysfunction, which exacerbates sarcopenia by interfering with other related degradation processes. Endurance exercise in old age increases the expression of Calcineurin and mTORC1, which contributes to TFE3 activation from the first session. TFE3 then improves lysosome biogenesis and other processes involved in waste product decomposition and cellular regeneration, minimizing sarcopenia and muscle wasting. The basal levels of Calcineurin gene expression and TFE3 protein activity increase with frequent endurance training and repeated stimulation of these factors. Endurance training regulates disruptive pathways by improving metabolism, which compensates for most of the muscle loss by inducing hypertrophy and a minor increase in muscle fibre number if any. Furthermore, while TFE3 activity increases with endurance exercise, it decreases over time after long-term training, indicating adaptation to training variables. Similarly, training adaptation reduces Calcineurin and mTORC1 post-session response, demonstrating the trainability of these three important factors for lysosome biogenesis and sarcopenia prevention, while emphasizing the importance of the training variety.

## Abbreviation list

A-SED: aged sedentary
A-SEDS: aged sedentary single session
A-L: Taged long-term trained
A-TS: aged trained-single session
E3=TFE3: transcription factor
MiT-TFE: microphthalmia family of transcription factors
MCOLN1: mucolipin-1
mTORC1: mechanistic target of rapamycin complex 1
ANOVA: analysis of variance

## Additional information

## Data availability statement

The data supporting the present findings are available from the corresponding author upon reasonable request.

## Competing interests

The authors declare that they have no competing interests.

## Author contributions

Based on CRediT taxonomy author contribution:

**B.A** and **H.R** contributed to Conceptualization. **B.A** contributed to Data curation, Formal analysis, Funding acquisition, Investigation, Visualization, Writing – original draft. **H.R** contributed to Project administration, Supervision. **B.A, H.R**, and **D.C** contributed to Validation, Writing – review and editing. All authors have read and approved the final version of this manuscript and agree to be accountable for all aspects of the work in ensuring that questions related to the accuracy or integrity of any part of the work are appropriately investigated and resolved. All persons designated as authors qualify for authorship, and all those who qualify for authorship are listed.

## Acknowledgements

The authors thank Dr Zohre Mazaheri Teyrani for her assistance in the ‘Resources’ part of this project.

## Funding

This research project was self-funded by Benyamin Askari as his master’s degree thesis at Kharazmi University.

## Notes

### Competing Interest Statement

The authors have declared no competing interest.

### Summary of Updates

1. there was a mistake in the abstract mentioning TFEB instead of TFE3. 2. there was duplicates in references and an uncounted citation due to misplening in discussion. 4. abbreviations are included. 5. abbreviated group names were inconsistent throughout the manuscript and are now corrected. 6.the introduction had redundant explainations on aging rate and humans statistics that are out of the scope of this project in aging biology. thses are removed. 7. all plot legends are updated.

## References

Baek, K.-W., Kim, S.-J., Kim, B.-G., Jung, Y.-K., Hah, Y.-S., Moon, H. Y., Yoo, J.-I., Park, J. S., & Kim, J.-S. (2022). Effects of lifelong spontaneous exercise on skeletal muscle and angiogenesis in super-aged mice. PLOS ONE, 17(8), e0263457. 10.1371/journal.pone.0263457

Bilski, J., Pierzchalski, P., Szczepanik, M., Bonior, J., & Zoladz, J. (2022). Multifactorial Mechanism of Sarcopenia and Sarcopenic Obesity. Role of Physical Exercise, Microbiota and Myokines. Cells, 11(1), 160. 10.3390/cells11010160

Carosi, J. M., Fourrier, C., Bensalem, J., & Sargeant, T. J. (2022). The mTOR–lysosome axis at the centre of ageing. FEBS Open Bio, 12(4), 739–757. 10.1002/2211-5463.13347

Charan, J., & Kantharia, N. D. (2013). How to calculate sample size in animal studies? Journal of Pharmacology and Pharmacotherapeutics, 4(4), 303–306. 10.4103/0976-500X.119726

Corrotte, M., & Castro-Gomes, T. (2019). Lysosomes and plasma membrane repair. In Current topics in membranes (1st ed., Vol. 84, pp. 1– 16). Elsevier Inc. 10.1016/bs.ctm.2019.08.001

Jorgenson, K. W., Phillips, S. M., & Hornberger, T. A. (2020). Identifying the Structural Adaptations that Drive the Mechanical Load-Induced Growth of Skeletal Muscle: A Scoping Review. Cells, 9(7), 1658. 10.3390/cells9071658

Kruschke, J. K. (2021). Bayesian Analysis Reporting Guidelines. Nature Human Behaviour, 5(10), 1282–1291. 10.1038/s41562-021-01177-7

Leary, S., Underwood, W., Anthony, R., Cartner, S., Grandin, T., Greenacre, C., Gwaltney-brant, S., Mccrackin, M. A., Meyer, R., Miller, D., Shearer, J., Turner, T., Equine, T., Yanong, R., Johnson, C. L., & Patterson-kane, E. (2020). AVMA Guidelines for the Euthanasia of Animals: 2020 Edition *. American Veterinary Medical Association.

Liang, J., Zeng, Z., Zhang, Y., & Chen, N. (2020). Regulatory role of exercise-induced autophagy for sarcopenia. Experimental Gerontology, 130(July 2019), 110789. 10.1016/j.exger.2019.110789

Martina, J. A., Diab, H. I., Brady, O. A., & Puertollano, R. (2016). TFEB and TFE3 are novel components of the integrated stress response. The EMBO Journal, 35(5), 479–495. 10.15252/embj.201593428

Memme, J. M., Erlich, A. T., Phukan, G., & Hood, D. A. (2021). Exercise and mitochondrial health. The Journal of Physiology, 599(3), 803–817. 10.1113/JP278853

National Research Council (US) Committee for the Update of the Guide for theCare and Use of Laboratory Animals. (2011). Guide for the Care and Use of Laboratory Animals (8th ed.). National Academies Press (US). http://www.ncbi.nlm.nih.gov/books/NBK54050/

Oliveira, A. N., & Hood, D. A. (2019). Exercise is mitochondrial medicine for muscle. Sports Medicine and Health Science, 1(1), 11–18. 10.1016/j.smhs.2019.08.008

Oliveira, A. N., Richards, B. J., Slavin, M., & Hood, D. A. (2021). Exercise Is Muscle Mitochondrial Medicine. Exercise and Sport Sciences Reviews, 49(2), 67–76. 10.1249/JES.0000000000000250

Quirós Cognuck, S., Reis, W. L., Silva, M., Debarba, L. K., Mecawi, A. S., de Paula, F. J. A., Rodrigues Franci, C., Elias, L. L. K., & Antunes-Rodrigues, J. (2020). Sex differences in body composition, metabolism-related hormones, and energy homeostasis during aging in Wistar rats. Physiological Reports, 8(20), e14597. 10.14814/phy2.14597

Roberts, F. L., & Markby, G. R. (2021). New Insights into Molecular Mechanisms Mediating Adaptation to Exercise; A Review Focusing on Mitochondrial Biogenesis, Mitochondrial Function, Mitophagy and Autophagy. Cells, 10(10), 2639. 10.3390/cells10102639

Tamaki, T., & Uchiyama, S. (1995). Absolute and relative growth of rat skeletal muscle. Physiology & Behavior, 57(5), 913–919. 10.1016/0031-9384(94)00359-D

Triolo, M., & Hood, D. A. (2021). Manifestations of Age on Autophagy, Mitophagy and Lysosomes in Skeletal Muscle. Cells, 10(5), 1054. 10.3390/cells10051054

Wang, C., Liang, J., Ren, Y., Huang, J., Jin, B., Wang, G., & Chen, N. (2022). A Preclinical Systematic Review of the Effects of Chronic Exercise on Autophagy-Related Proteins in Aging Skeletal Muscle. Frontiers in Physiology, 13(July), 930185. 10.3389/fphys.2022.930185

Wasserstein, R. L., Schirm, A. L., & Lazar, N. A. (2019). Moving to a World Beyond “p < 0.05”. The American Statistician, 73(Sup1), 1– 19. 10.1080/00031305.2019.1583913

Wróbel, M., Cendrowski, J., Szymańska, E., Grębowicz-Maciukiewicz, M., Budick-Harmelin, N., Macias, M., Szybińska, A., Mazur, M., Kolmus, K., Goryca, K., Dąbrowska, M., Paziewska, A., Mikula, M., & Miăczyńska, M. (2022). ESCRT-I fuels lysosomal degradation to restrict TFEB/TFE3 signaling via the Rag-mTORC1 pathway. Life Science Alliance, 5(7), e202101239. 10.26508/lsa.202101239

Yang, Y., Xu, M., Zhu, X., Yao, J., Shen, B., & Dong, X.-P. (2019). Lysosomal Ca2+ release channel TRPML1 regulates lysosome size by promoting mTORC1 activity. European Journal of Cell Biology, 98(2–4), 116–123. 10.1016/j.ejcb.2019.05.001

Zhu, L., Ye, T., Tang, Q., Wang, Y., Wu, X., Li, H., & Jiang, Y. (2016). Exercise Preconditioning Regulates the Toll-Like Receptor 4/Nuclear Factor-κB Signaling Pathway and Reduces Cerebral Ischemia/Reperfusion Inflammatory Injury: A Study in Rats. Journal of Stroke and Cerebrovascular Diseases, 25(11), 2770–2779. 10.1016/j.jstrokecerebrovasdis.2016.07.033

